# Order of amino acid recruitment into the genetic code resolved by Last Universal Common Ancestor’s protein domains

**DOI:** 10.1101/2024.04.13.589375

**Authors:** Sawsan Wehbi, Andrew Wheeler, Benoit Morel, Nandini Manepalli, Bui Quang Minh, Dante S. Lauretta, Joanna Masel

## Abstract

The current “consensus” order in which amino acids were added to the genetic code is based on potentially biased criteria, such as absence of sulfur-containing amino acids from the Urey-Miller experiment which lacked sulfur. More broadly, abiotic abundance might not reflect biotic abundance in the organisms in which the genetic code evolved. Here, we instead identify which protein domains date to the last universal common ancestor (LUCA), then infer the order of recruitment from deviations of their ancestrally reconstructed amino acid frequencies from the still-ancient post-LUCA controls. We find that smaller amino acids were added to the code earlier, with no additional predictive power in the previous “consensus” order. Metal-binding (cysteine and histidine) and sulfur-containing (cysteine and methionine) amino acids were added to the genetic code much earlier than previously thought. Methionine and histidine were added to the code earlier than expected from their molecular weights, and glutamine later. Early methionine availability is compatible with inferred early use of S-adenosylmethionine, and early histidine with its purine-like structure and the demand for metal-binding. Even more ancient protein sequences — those that had already diversified into multiple distinct copies prior to LUCA — have significantly higher frequencies of aromatic amino acids (tryptophan, tyrosine, phenylalanine and histidine), and lower frequencies of valine and glutamic acid than single copy LUCA sequences. If at least some of these sequences predate the current code, then their distinct enrichment patterns provide hints about earlier, alternative genetic codes.

**Significance Statement:** The order in which the amino acids were added to the genetic code was previously inferred from consensus among forty metrics. Many of these reflect abiotic abundance on ancient Earth. However, the abundances that matter are those within primitive cells that already had sophisticated RNA and perhaps peptide metabolism. Here, we directly infer the order of recruitment from the relative ancestral amino acid frequencies of ancient protein sequences. Small size predicts ancient amino acid enrichment better than the previous consensus metric does. We place metal-binding and sulfur-containing amino acids earlier than previously thought, highlighting the importance of metal-dependent catalysis and sulfur metabolism to ancient life. Understanding early life has implications for our search for life elsewhere in the universe.

## Introduction

The modern genetic code was likely assembled in stages, hypothesized to begin with “early” amino acids present on Earth before the emergence of life (possibly delivered by extraterrestrial sources such as asteroids or comets), and ending with “late” amino acids requiring biotic synthesis (1, 2). For example, the Urey-Miller experiment (3) has been used to identify which amino acids were available abiotically and are thus likely to have come earlier than those requiring biotic synthesis. The order of amino acid recruitment, from early to late, was inferred by taking statistical consensus among 40 different rankings (4), none of which constitute strong evidence on their own. On the basis of this ordering, Moosmann (5) hypothesized that the first amino acids recruited into the genetic code were those that were useful for membrane anchoring, then those useful for halophilic folding, then for mesophilic folding, then for metal binding, and finally for their antioxidant properties. However, a late role for metal-binding amino acids is puzzling; many metalloproteins date back to the Last Universal Common Ancestor’s (LUCA)’s proteome, where they are presumed to be key to the emergence of biological catalysis (6).

Indeed, the late status of some amino acids is disputed (7). For example, the Urey-Miller experiment (3) did not include sulfur, and so should not have been used to infer that the sulfur-containing amino acids cysteine and methionine were late additions. Methionine and homocysteine (a product of cysteine degradation) were detected in hydrogen sulfide (H_2_S)-rich spark discharge experiments, suggesting that methionine and cysteine could be abiotically produced (8). A nitrile-activated dehydroalanine pathway can produce cysteine from abiotic serine that is produced from a Strecker reaction (9), further demonstrating the possibility of its early chemical availability.

Histidine’s classification as abiotically unavailable also contributed to its annotation as late (4). While histidine can be abiotically synthesized from erythrose reacting with formamidine followed by a Strecker synthesis reaction (10), the reactant concentrations might have been insufficient in a primitive earth environment (11). More importantly, because histidine resembles a purine, even if histidine were abiotically unavailable, it might have had cellular availability at the time of genetic code construction (12), in an organism that biotically synthesized ribosomes, and that might also have already utilized amino acids and peptides. Indeed, histidine is the most commonly conserved residue in the active site of enzymes (13).

To directly infer the order of recruitment from protein sequence data, without reference to abiotic availability arguments, we consider that some of LUCA’s proteins were born prior to the completion of the genetic code (14). We predict that ancestrally reconstructed sequences from this era will be enriched in early amino acids and depleted in late amino acids. Previous analyses relied on conserved residues within a small number of LUCA proteins (15, 16). Here, we classify a larger set of protein-coding domains that date back to LUCA, rather than being more recently born, e.g., *de novo* from non-coding sequences or alternative reading frames (17, 18). We compare reconstructed ancient amino acid frequencies of the most ancient vs. moderately ancient protein cohorts, to deduce the order in which amino acids were incorporated into the genetic code.

We take advantage of gene-tree species-tree reconciliation methods (19) to infer LUCA’s protein sequences. Previous analyses focused on the age of orthologous gene families (20-22); ours is the first to infer which protein domains date back to LUCA. Protein domains are the basic units of proteins, that can fold, function, and evolve independently (23). Proteins often contain multiple protein domains, each of which might have a different age (Figure 1). For the purpose of inferring ancient amino acid usage, what matters is the age of the protein domain, not that of the whole protein that it is part of. We use protein domain annotations from the Pfam database (24). We recognize Pfams present in LUCA by trimming horizontal gene transfer (HGT) events, and by exploiting long archaeal-bacterial branches (Figure 2; see Methods for details).

**Figure 1.**
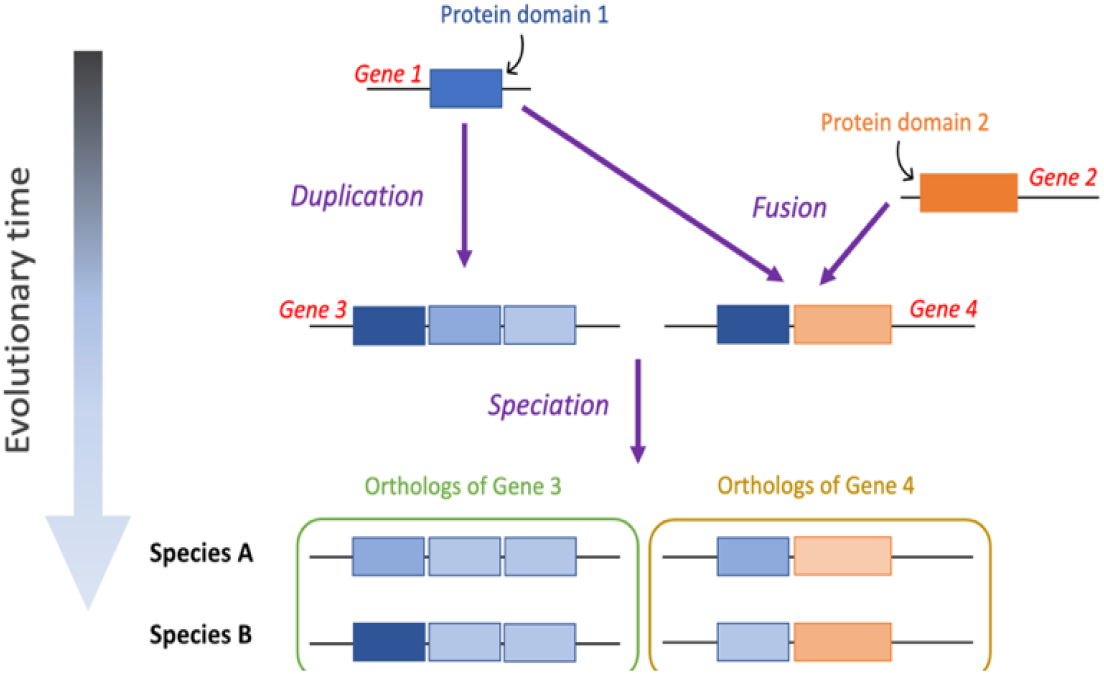
The evolutionary history of a protein domain may date back further in time than that of the whole-gene ortholog that it is part of. Multi-domain genes 3 and 4 originated around the same time. However, they are made up of two protein domains (blue & orange boxes) that emerged and diverged at different points in time – domain 1 is older than domain 2.

**Figure 2.**
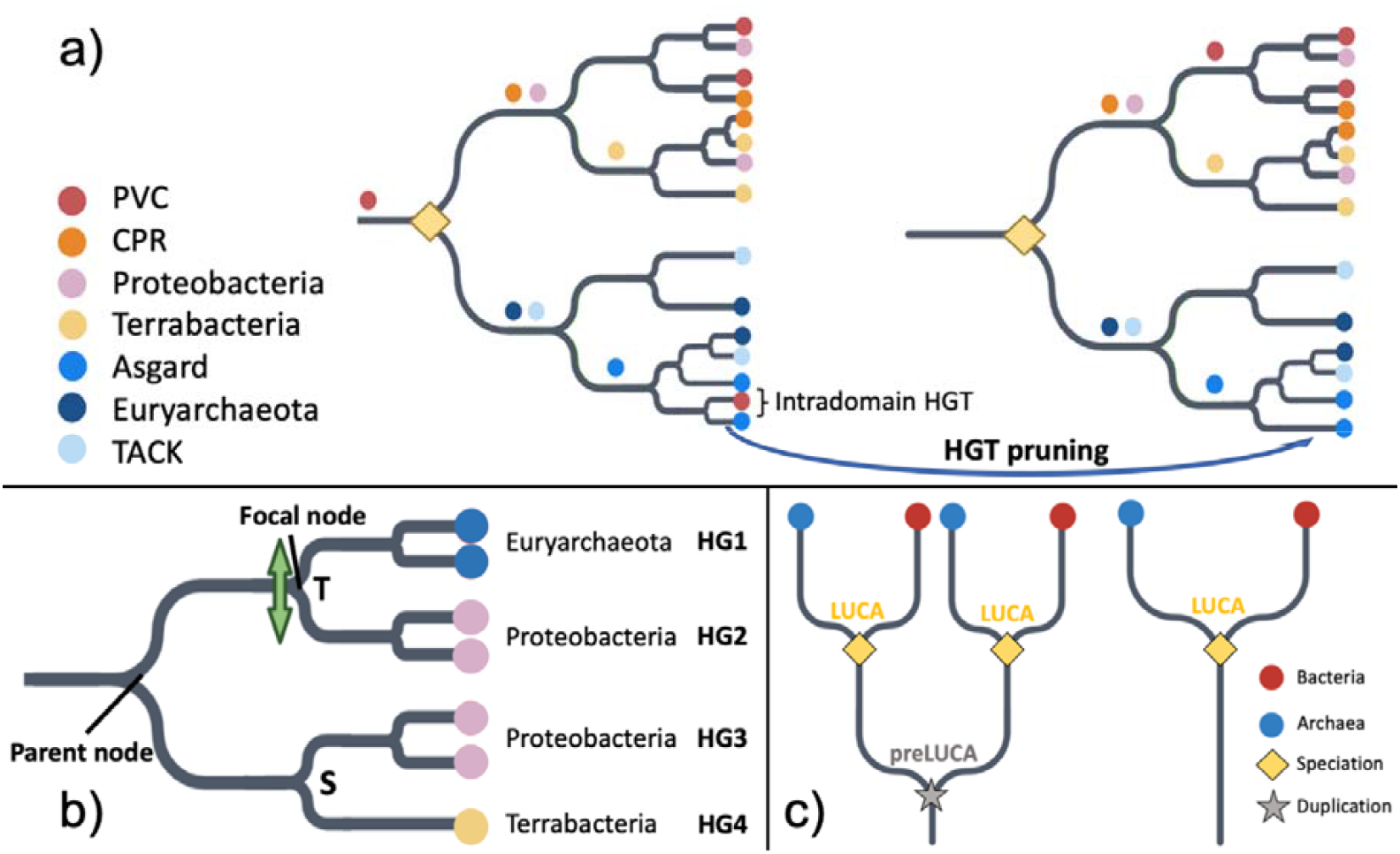
Criteria for (a) LUCA Pfam annotation, (b) identifying HGT to be filtered, and (c) pre-LUCA Pfam annotation. Details are in Methods, with a brief summary here. a) Pruning HGT between archaea and bacteria reveals a LUCA node as dividing bacteria and archaea at the root. Colored circles are indicated just upstream of the most recent common ancestor (MRCA) of all copies of that Pfam found within the same taxonomic supergroup. We recognize a total of five bacterial supergroups (FCB, PVC, CPR, Terrabacteria and Proteobacteria (75, 76)) and four archaeal supergroups (TACK, DPANN, Asgard and Euryarchaeota (77, 78)); only 4 out of 5 bacterial supergroups and 3 out of 4 archaeal supergroups are shown. The yellow diamond indicates LUCA as a speciation event between archaea and bacteria. We do not assume that the LUCA coalescence timing was the same for every Pfam (94). Prior to HGT pruning, PVC sequences can be found on either side of the two lineages divided by the root. After pruning intradomain HGT, four MRCAs are found one node away from the root, and three more MRCAs are found two nodes away from the root, fulfilling our other LUCA criterion described in the Methods, namely presence of at least three bacterial and at least two archaeal supergroup MRCAs one to two nodes away from the root. b) Criteria for pruning likely HGT between archaea and bacteria (see Methods for details). We partition into monophyletic groups of sequences in the same supergroup; in this example, there are four such groups, representing two bacterial supergroups and one archaeal supergroup. There is one ‘mixed’ node, separating an archaeal group (HG1) from a bacterial group (HG2). It is also annotated by GeneRax (19) as a transfer ‘T’. The bacterial nature of groups 3 and 4 indicates a putative HGT direction from group 2 to group 1. Group 2 does not contain any Euryarchaeota sequences, meeting the third and final requirement for pruning of group 1. If neither Proteobacteria or Euryarchaeota sequences were present among the other descendants of the parent node, both groups 1 and 2 would be considered acceptors of a transferred Pfam and would both be pruned from the tree. c) Pre-LUCA Pfams have at least two nodes annotated as LUCA.

## Results

### Ancient protein domain classifications agree with whole-gene classifications

We classify 969 Pfams and 445 clans (sets of one or more Pfams that are evolutionary related) as present in LUCA (Figures 3a and 3b; detailed lists in Supplementary Tables 1 and 2). We compare these to the 3055 Pfams and 1232 clans that we classify as ancient but post-LUCA (including Last Bacterial Common Ancestor (LBCA) and Last Archaeal Common Ancestor (LACA) candidates). Encouragingly, 88.6% of Pfams that we annotate as pre-LUCA or LUCA are contained within genes annotated by Moody et al. (21) as present in LUCA with more than 50% confidence, when present in their dataset (Figure 3c). This level of agreement far exceeds earlier works (22).

**Figure 3.**
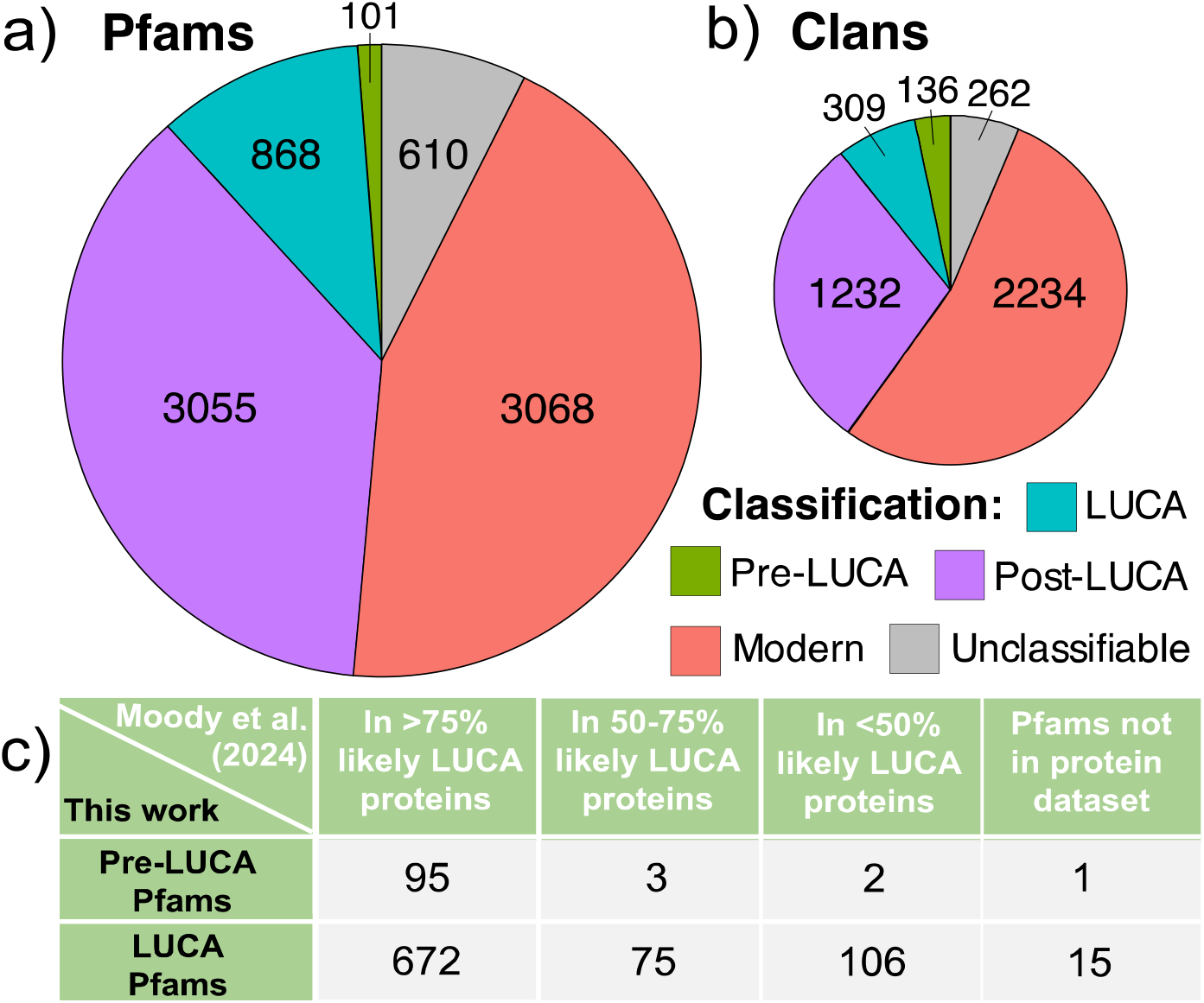
Pfams (a) and clans (b) classified as ancient are well validated by the whole gene annotations of Moody et al. (21) (c). a) Ancient post-LUCA Pfam classifications include 285 LACA candidates and 2770 LBCA candidates (more analysis would be required to rule out extensive HGT within archaea or bacteria). Modern Pfams are distributed among the prokaryotic supergroups as follows: 9 CPR, 210 FCB, 942 Proteobacteria, 51 PVC, 1111 Terrabacteria, 2 Asgard, 49 TACK, and 177 Euryarchaeota. In addition to supergroup-specific modern Pfams, we classified another 1097 Pfams, present in exactly two bacterial supergroups, as modern post-LBCA. We deemed 15 Pfams unclassifiable due to high inferred HGT rates, 397 due to uncertainty in rooting, and 198 due to ancient rooting combined with absence from too many supergroups (see Methods). b) Pre-LUCA clans contain at least two LUCA-classified Pfams or one pre-LUCA Pfam, whereas LUCA clans contain exactly one LUCA Pfam. Ancient post-LUCA clans contain no LUCA, pre-LUCA, or unclassified Pfams; they include an ancient post-LUCA Pfam or at least two modern Pfams covering at least two supergroups from only one of either bacteria or archaea. Modern clans include Pfams whose root is assigned at the origin of one supergroup. Finally, unclassifiable clans did not meet any of our clan classification criteria, e.g., because they included both post-LUCA and unclassifiable Pfams. c) 98% of our pre-LUCA Pfams and 87% of our LUCA Pfams are present in genes annotated by as present in LUCA with more than 50% confidence, when present in their dataset. We mapped all Clusters of Orthologous Genes (COGs) (95) in the Moody et al. (21) supplementary dataset (STable_1.csv) to UniProt IDs (96) using the EggNOG 5.0 database (97). We then identified their associated Pfams using the ‘Pfam-A.regions.uniprot.tsv’ file downloaded from the Pfam FTP site (https://pfam-docs.readthedocs.io/en/latest/ftp-site.html#current-release) (24) on May 28^th^, 2024. Our protein to Pfam ID mappings are available in ‘Protein2Domain_mappings’ at https://doi.org/10.6084/m9.figshare.27191274.v1.

In agreement with the Moody et al. (21) classification of LUCA metabolism, almost all Pfams associated with enzymes in hydrogen metabolism, assimilatory nitrate and sulfate reduction pathways, and the Wood-Ljungdahl pathway date back to LUCA (Supplementary Table 3). Our results also support a, post-LUCA, bacterial origin of nitrogen fixation (21, 25) (Supplementary Table 3). We assign to LUCA the complete set of amino acid-tRNA synthetase-associated anti-codon binding domains found in modern prokaryotes. Here, focusing on complete genes would have been problematic, because accessory amino acid-tRNA synthetase-associated domains (e.g. PF04073 and PF13603, which deacylate misacylated tRNA) were sometimes added later.

We also checked the antiquity of the cofactor/cosubstrate S-adenosylmethionine (SAM) (26), both with respect to SAM biosynthesis and SAM usage. In agreement with past work attributing the SAM biosynthesis enzyme methionine adenosyltransferase to LUCA (27, 28), we assign its single Pfam (PF01941) to LUCA (the corresponding COG1812 is not analyzed by Moody et al. (21)). In agreement with past work attributing SAM-dependent methyltransferases to LUCA (29), Moody et al. (21) assign the RsmB/RsmF family (COG0144), which methylates 16S rRNA, more than 75% confidence of being present in LUCA, and we also classify its SAM-binding Rossman fold Pfam (PF01189) as LUCA. In agreement with (30, 31), Moody et al. (21) assign the SAM-binding tRNA methylthiolase (COG0621) to LUCA with more than 75% confidence, and we confirm the pre-LUCA status of its associated Radical SAM, TIM-barrel-related Pfam (PF04055). In agreement with attribution of polyamines to LUCA (32) we assign to LUCA the one Pfam (PF02675) of S-adenosylmethionine decarboxylase, which acts on SAM in the first step of polyamine synthesis; the antiquity of corresponding COG1586 is not further confirmed by Moody et al. (21).

### Hydrophobic amino acids are more interspersed within ancient proteins

Interspersion of hydrophobic amino acids away from one another along the primary sequence is believed to mitigate risks from protein misfolding, while still enabling correct folding (33-35). Older sequences have previously been found to have greater interspersion among their hydrophobic residues, indicating more sophisticated protein folding (14, 36), likely due to survivorship bias (37). Our Pfam age classifications confirm the antiquity of this trend, previously observed only for animal sequences. LUCA Pfams show even more hydrophobic interspersion than the still-ancient ‘post-LUCA’ Pfams that include LACA candidates and LBCA candidates (Supplementary Figure 1; Wilcoxon rank sum test; p = 0.02). Post-LUCA Pfams in turn have more hydrophobic interspersion than ‘modern’ Pfams that are specific to particular prokaryotic supergroups (Wilcoxon rank sum test; p = 0.02).

### LUCA’s protein sequences were depleted in larger amino acids

Clans present in LUCA were born before the divergence of Archaea and Bacteria, some potentially prior to the completion of the genetic code. If newly recruited amino acids were added slowly, the contemporary descendants of LUCA clans will show signs of ancestral depletion in amino acids that were added late to the genetic code. We first focus on clans present in one copy in LUCA (denoted “LUCA clans”), excluding those that had already duplicated and diverged into multiple surviving lineages (denoted “pre-LUCA clans”). We score ancestral amino acid enrichment and depletion as relative to still-ancient post-LUCA clans, which represent amino acid usage from the standard genetic code of all 20 amino acids, plus any ascertainment biases. This ratio, reflecting ancient amino acid usage, is not confounded with the effects of temperature, pH, oxygen tolerance, salinity, GC content, or transmembrane status on amino acid frequencies (Supplementary Figures 2a-f). Indeed, LUCA usage is similar in the very different biophysical context of a transmembrane site (Supplementary Figure 3).

Smaller amino acids are enriched in LUCA (Figure 4a; weighted *R*^2^ = 0.48, *p* = 0.0005). Results are similar using a restricted set of Pfams validated by Moody et al. (21) (weighted *R*^2^ = 0.44, *p* = 0.001). As a negative control for methodological artifacts, the ancestral amino acid usage of post-LUCA clans relative to modern clans is not correlated with molecular weight (*p* = 0.9).

**Figure 4.**
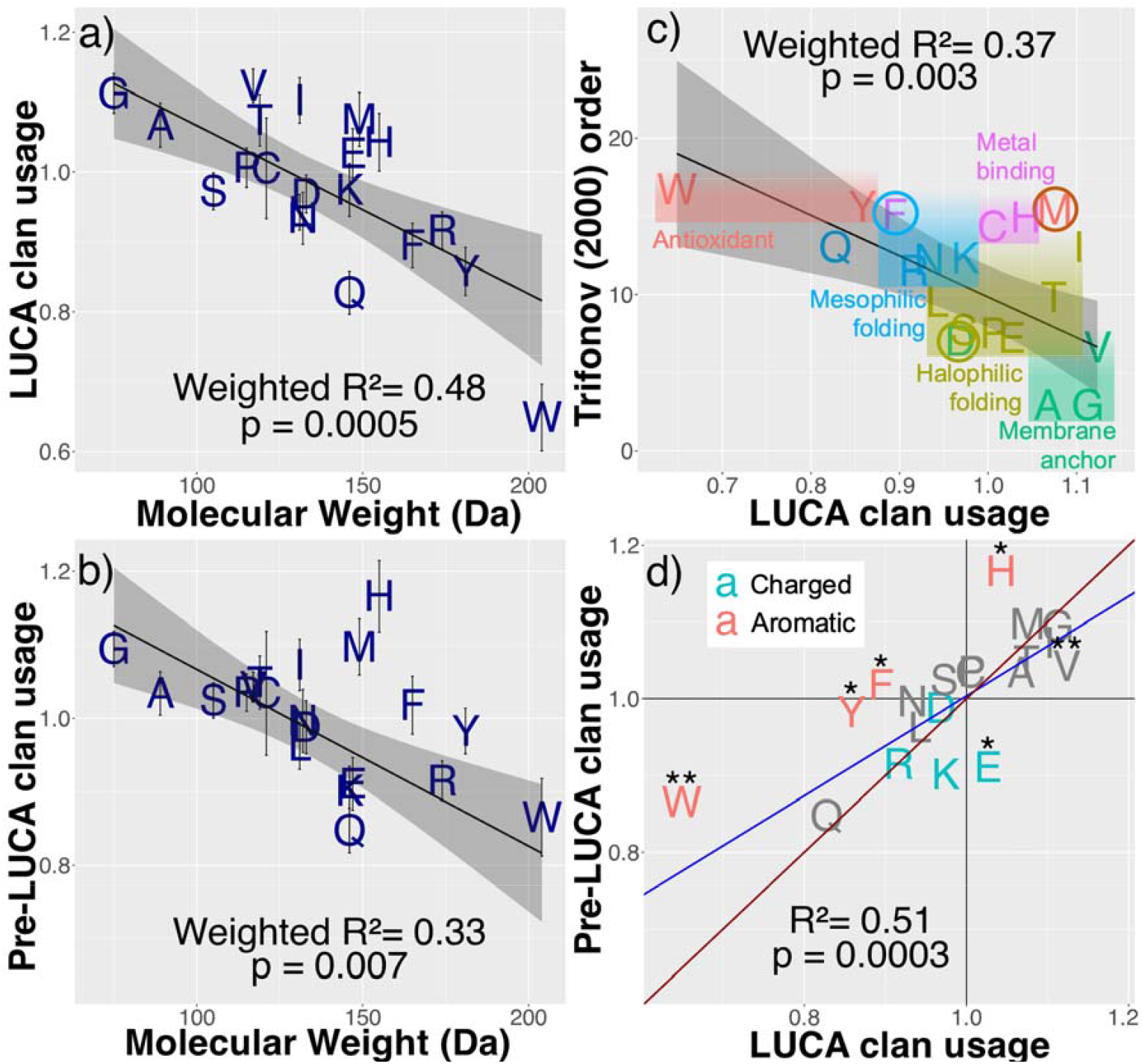
LUCA is enriched for smaller amino acids, with subtle differences between single copy LUCA vs. multi-copy pre-LUCA sequences. Ancestrally reconstructed amino acid frequencies in LUCA and pre-LUCA clans are shown relative to those in ancient post-LUCA clans. a) LUCA clans and b) pre-LUCA clans are enriched for amino acids of smaller molecular weight. Weighted model 1 regression lines are shown in black with 95% confidence interval grey shading. Error bars indicate standard errors. c) Character colors show the assignments of Moosmann (5); colored circles indicate our re-assignments. We reclassify phenylalanine (F) because it is enriched in proteins in mesophiles compared to their orthologs in thermophiles and hyperthermophiles (98). We reclassify aspartic acid (D) because the surfaces of proteins within halophilic bacteria are highly enriched in aspartic acid compared to in the surfaces of non-halophilic mesophilic and thermophilic bacteria, in a manner that cannot be accounted for by the dinucleotide composition of the halophilic genomes (99). The brown circle around methionine highlights that while it might not be utilized against reactive oxygen species, it might once have been against ancient reactive sulfur species. d) Model 2 Deming regression (accounting for standard errors in both variables, implemented in deming() version 1.4-1 (100)) in blue shows that pre-LUCA enrichments are not more extreme versions of LUCA enrichments, lying on the wrong side of the y=x red line. We include the imidazole-ring-containing H as aromatic. Asterisks (*) indicate statistically different amino acid frequencies between pre-LUCA and LUCA (Welch two sample t-test, p<0.05 and p<0.01).

### Revised Order of Amino Acid Recruitment

Figure 4c visualizes how LUCA’s amino acid enrichments compare to Trifonov’s consensus order (4). While they are correlated (weighted *R*^2^ = 0.37, *p* = 0.003), this association disappears in a weighted multiple regression with both molecular weight (*p* = 0.03) and Trifonov’s (4) order (*p* = 0.9) as predictors (weighted *R*^2^ = 0.48). This is also true using Trifonov’s revised 2004 order based on 60 metrics (38) (weighted *R*^2^ = 0.34, *p* = 0.006 on its own; *p* = 0.9 when molecular weight is also a predictor of LUCA usage). This suggests that some of Trifonov’s 40-60 metrics made his estimates of the order of recruitment worse rather than better. We use enrichment in LUCA to re-classify VGIMTAHEPC as ‘early’ and depletion to classify KSDLNRFYQW as ‘late’. More precise estimation of the order of recruitment, with standard errors, is given in Table 1.

**Table 1.** LUCA and pre-LUCA clans’ ancestral amino acid frequencies are divided by post-LUCA clan’s ancestral amino acid frequencies to produce measures of relative usage. The standard errors of the amino acid usages were calculated using an approximation derived from a Taylor expansion of the ratio (90). For each of the 20 ancestral amino acid frequencies, the standard errors of the weighted means across all the clans within the LUCA and pre-LUCA phylostrata (weighted by the maximum number of ancestral sites across all Pfams in a given clan) were calculated using the weighted_se() function in the diagis R package (89)(See Methods for more detail).

| Amino acid | LUCA usage | LUCA usage<br>standard<br>error | Pre-LUCA usage | Pre-LUCA usage<br>standard error |
| --- | --- | --- | --- | --- |
| V | 1.12 | 0.0241 | 1.04 | 0.0205 |
| G | 1.11 | 0.0283 | 1.09 | 0.0241 |
| I | 1.1 | 0.0325 | 1.07 | 0.0351 |
| M | 1.08 | 0.0386 | 1.1 | 0.0383 |
| A | 1.07 | 0.0317 | 1.03 | 0.0297 |
| T | 1.07 | 0.0369 | 1.05 | 0.0362 |
| H | 1.04 | 0.0416 | 1.17 | 0.0486 |
| E | 1.03 | 0.0357 | 0.911 | 0.0357 |
| C | 1.01 | 0.0722 | 1.03 | 0.0844 |
| P | 1.01 | 0.0282 | 1.04 | 0.0255 |
| K | 0.974 | 0.038 | 0.901 | 0.0334 |
| S | 0.972 | 0.0265 | 1.02 | 0.0239 |
| D | 0.968 | 0.027 | 0.988 | 0.0363 |
| L | 0.942 | 0.0256 | 0.962 | 0.032 |
| N | 0.934 | 0.0374 | 0.996 | 0.0432 |
| R | 0.916 | 0.0265 | 0.915 | 0.0271 |
| F | 0.895 | 0.032 | 1.02 | 0.0394 |
| Y | 0.858 | 0.0341 | 0.982 | 0.0309 |
| Q | 0.827 | 0.031 | 0.847 | 0.0304 |
| W | 0.649 | 0.0476 | 0.865 | 0.0526 |

We place glutamine (Q or Gln) as the second last amino acid, much later than Trifonov (4) inferred. Consistent with its late addition, Gln-tRNA synthetase (GlnRS) is either absent in prokaryotes, or acquired via horizontal gene transfer from eukaryotes (39). Prokaryotes that lack GlnRS perform tRNA-dependent amidation of Glu mischarged to Gln-tRNA by GluRS, forming Gln-acylated Gln-tRNA via amidotransferase. The core catalytic domain (PF00587), shared between the GlnRS and GluRS paralogs, is present in LUCA and can indiscriminately acylate both Gln-tRNA and Glu-tRNAs with Glu (40).

### Metal-binding and sulfur-containing amino acids were added early to the genetic code

Methionine (M), cysteine (C), and histidine (H) are all enriched in LUCA, despite previous annotation as late additions to the genetic code (Figure 4c). C and H are the most frequently used amino acids for binding iron, zinc, copper, and molybdenum, and H, aspartic acid (D) and glutamic acid (E or Glu) for binding manganese and cobalt (Figure 2D of (41)). Binding can either be to a metal ion, or to iron-sulfur (FeS) clusters, usually via C but sometimes via H or D (42). Binding these transition metals is key to catalysis (43). Figure 4a is incompatible with C, H, D, or E being late additions, and indeed H is more enriched than one would expect from its molecular weight.

C and M are the only sulfur-containing amino acids in the contemporary genetic code. Contemporary prokaryotes living in H_2_S-rich environments use more C and M than matched species (Supplementary Figure 4); LUCA’s C and M enrichment might thus reflect an environment rich in H_2_S.

Moosmann (5) classified M, tryptophan (W), and tyrosine (Y) as antioxidants, because he believed them to protect the overall protein structure from oxidative stress via sacrificial oxidization. For instance, surface M residues can be reversibly oxidized to form methionine sulfoxide (44). This might have driven isoleucine recoding to methionine in mitochondria (45, 46). However, proteins in aerobes are enriched in W and Y but not in M (47). Our results also separate early M from late Y and W (Figure 4). We speculate that methionine, abundant due to early life’s use of SAM, might have protected against reactive sulfur species such as sulfide (S^2-^), which were present in early, H_2_S-rich environments (48). Our results are then partially compatible with Granold et al.’s (49) view that Y and W (but not M) were added to complete the modern genetic code after reactive oxygen species became the main oxidizing threat.

### Pre-LUCA clans hint at more ancient genetic codes

We expected pre-LUCA enrichments and depletions to be more extreme than for LUCA, but only H fits this prediction (Figure 4d), with significantly higher frequencies in pre-LUCA than in LUCA.

There is nevertheless a strong overall correlation between LUCA and pre-LUCA usages (*R*^2^ = 0.51, *p* = 0.0003). Pre-LUCA, like LUCA, is strongly depleted in Q, supporting the inference that Q, not Y, was the 19^th^ amino acid recruited into the standard genetic code. Pre-LUCA usage does not correlate with Trifonov’s consensus order (4) (*p* = 0.2), and correlates more weakly with molecular weight (Figure 4b) (weighted *R*^2^ = 0.33, *p* = 0.007).

H is one of six amino acids with significantly different frequencies in pre-LUCA vs. LUCA. All three of the canonical, benzene-ring bearing, aromatic amino acids (W, Y, and phenylalanine (F)), as well as the imidazole-ring containing H, are more common in pre-LUCA than in LUCA (Figure 4d, Welch 2-sample t-test; *p* = 0.03, 0.001, 0.03 and 0.01, respectively; 2.4% vs 2.1% H, 1.2% vs 0.9% W, 3.1% vs. 2.8% Y, and 4.1% vs. 3.7% F). Glutamic acid (E) and Valine (V) are less common in pre-LUCA than in LUCA (Welch 2-sample t-test; *p* = 0.01 and 0.004, respectively; 7.3% vs. 8.2% E, 7.5% vs. 8.1% V).

More W in pre-LUCA than LUCA is particularly surprising, because there is scientific consensus that W was the last of the 20 canonical amino acids to be added to the genetic code. Therefore, we manually inspected the pre-LUCA Pfam with the highest tryptophan frequency (3.1%): PF00133, the core catalytic domain of the tRNA synthetases of leucine (L), isoleucine (I), and valine (V). Each of these three synthetases has well-separated archaeal and bacterial branches, confirming its pre-LUCA dating (Supplementary Figure 5). Highly conserved tryptophan sites regulate the size of the amino acid binding pocket, allowing the synthetases to discriminate among I, L, and V (50). There are also conserved I and V sites in the common ancestor of the I and V tRNA synthetases, indicating that discrimination between the two happened prior to the evolution of the synthetases currently responsible for the discrimination (51). This suggests that an alternative, more ancient system predated the modern genetic code, and in particular predated the evolution of super-specific, cognate aaRSs (51).

## Discussion

The evolution of the current genetic code proceeded via stepwise incorporation of amino acids, driven in part by changes in early life’s environment and requirements. Contemporary proteins retain information about which amino acids were part of the code at the time of their birth, allowing us to infer the order of recruitment on the basis of enrichment or depletion in LUCA’s protein domains. Smaller amino acids were added to the code first, and when this is accounted for, there is no further information in Trifonov’s (4) widely used ‘consensus’ order based on 40 metrics, some of dubious relevance. The sulfur-containing amino acids C and M were incorporated earlier than previously thought, likely because those metrics included experiments conducted in the absence of sulfur. Q was added later than previously thought, in agreement with evidence from glutamyl-tRNA synthetases. M and H were added to the code earlier than expected from their molecular weights, and Q later. Even more ancient amino acid usage, in sequences that had already duplicated and diverged pre-LUCA, shows significantly higher frequencies of the aromatic amino acids W, Y, F, and H, and significantly lower frequencies of E and V.

If LUCA lived in a H_2_S-rich environment (48, 52), M residues could have protected proteins against sulfur-mediated oxidative stress. M would furthermore have had high biotic availability as the precursor (53) and product (54) of SAM, given our finding that LUCA made and used SAM. The potentially sulfur-rich nature of early terrestrial life is context for astrobiology investigations of sulfur-rich environments on Mars and Europa, with associated biosignatures key to life detection (55).

An early role for H is compatible with a key role for metal binding in early life. It also resolves the previous puzzle that the ancestral, conserved region of all Class I aaRSs contains a histidine-rich HIGH motif (56, 57). The lack of abiotic availability was key to H’s previous annotation as late, but biotic availability of H in an RNA-dominant biotic context would have been sufficient. The importance of abiotic availability (58, 59) to the origins of the genetic code remains unclear. We note that ongoing research on plausible prebiotic syntheses in cyanosulfidic environments (60) and alkaline hydrothermal vents (61) is reshaping our understanding of which amino acids were accessible to early life. Amino acid abundances obtained from asteroid sample returns will also soon contribute (62).

Our results offer an improved approximation of the order of recruitment of the twenty amino acids into the genetic code under which contemporary protein-coding sequences were born. This order need not match the importance or abundance with which amino acids were used by still earlier life forms, nor during the prebiotic to biotic transition. Instead of using Trifonov’s assignments (4) to capture the order in which amino acids were recruited into our genetic code, we recommend using the LUCA amino acid enrichment values plotted on the y-axis of Figure 4a, which can be found together with their standard errors in Table 1.

More broadly, coding for different amino acids might have emerged at similar times but in different biogeochemical environments. The temporal order of recruitment that we infer based on LUCA sequences is not the temporal order for coding as a whole, but for the ancestor of the modern translation machinery. Indeed, horizontal gene transfer of the tRNAs coupled with their cognate aminoacyl tRNA synthetases might have brought the diverse components of the modern translation machinery together (63). This further emphasizes that the time of origin of the translation machinery’s components need not match the time of their incorporation into the surviving ancestral lineage.

The construction of the genetic code was tethered to the evolution of the ribosome (64). If the ribosome’s exit tunnel, whose formation and subsequent extension was key to ribosome evolution (65), limited the size of the amino acids passing through, its progressive dilation could explain the strong relationship between amino acid size and order of recruitment evidenced in LUCA clans. If older, alternative codes were not similarly limited, this would explain why amino acid size is a weaker predictor of pre-LUCA’s amino acid usage compared to LUCA’s amino acid usage.

To explain the different enrichments of pre-LUCA versus LUCA sequences, as well as the surprising conservation of some sites prior to the emergence of the aaRSs that distinguish the relevant amino acids, we propose that some pre-LUCA sequences are older than the current genetic code, perhaps even tracing back to a peptide world at the dawn of precellular life (7). Stepwise construction of the current code and competition among ancient codes could have occurred simultaneously (66, 67). Ancient codes might also have used non-canonical amino acids, such as norvaline and norleucine (68) which can be recognized by LeuRS (69, 70). Along with having different genetic codes, we speculate that pre-LUCA and LUCA might have existed in different geochemical settings. For instance, if pre-LUCA ancestors inhabited alkaline hydrothermal vents, where abiotically produced aromatic amino acids have been found (61), this would explain their enrichment in pre-LUCA relative to LUCA. We note that abiotic synthesis of aromatic amino acids might be possible in the water-rock interface of Enceladus’s subsurface ocean, which is speculated to be analogous to terrestrial alkaline hydrothermal vents (71).

Perhaps the biggest mystery is how sequences such as the common ancestor of L/I/V-tRNA synthetase, which were translated via alternative or incomplete genetic codes, ended up being re-coded for translation by the direct ancestor of the canonical genetic code. Harmonization of genetic codes facilitated innovation sharing via HGT, making it advantageous to use the most common code, driving code convergence (72, 73). Only once a common code was established did HGT drop to levels such that a species tree became apparent, i.e. the LUCA coalescence point corresponds to convergence on a code (72). Our identification of pre-LUCA sequences provides a rare source of data about early, alternative codes.

## Materials and Method

### Pfam sequences

We downloaded genomes of 3562 prokaryotic species from NCBI that were present in the Web of Life (WoL): Reference phylogeny of microbes (74) in August 2022. We classified them into five bacterial supergroups (FCB, PVC, CPR, Terrabacteria and Proteobacteria (75, 76)) and four archaeal supergroups (TACK, DPANN, Asgard and Euryarchaeota (77, 78)). We included incomplete genomes, to enhance coverage of underrepresented supergroups.

We assign ages not to whole proteins but to each of their protein domain constituents. We used InterProScan (79) to identify instances of each Pfam domain (24). We excluded Pfams with fewer than 50 instances across all downloaded genomes. We also excluded 9 Pfams marked “obsolete” starting July 2023. Among the remaining 8282 Pfams, 2496 Pfams had more than 1000 instances. We downsampled these to balance representation across the two taxonomic domains (archaea and bacteria). For instance, a Pfam with 2000 bacterial and 500 archaeal instances was downsampled by retaining all 500 archaeal sequences plus a subset (randomly sampled without replacement) of 500 bacterial sequences.

The Pfam database includes annotations of “clans” of Pfams that share a common ancestor despite limited sequence similarity; for many analyses, we used clans rather than Pfams to ensure independent datapoints. We treated Pfams that were not annotated as part of a clan as single-entry clans, with clan ID equal to their Pfam ID.

### Pfam trees

We aligned downsampled sequences for each Pfam using MAFFT v.7 (80), to infer a preliminary tree with IQ-Tree (81), using a time non-reversible amino acid substitution matrix trained on the Pfam database (NQ.PFAM) (82), and no rate heterogeneity among sites. Because most Pfams are too short for reliable tree inference, we next reconciled preliminary Pfam trees with the WoL prokaryotic species tree (74) using GeneRax (19). While there is no perfect species tree for prokaryotes, reconciliation even with a roughly approximate tree can still provide benefits. We ran GeneRax twice. The first run used the LG amino acid substitution model, a gamma distribution with four discrete rate categories, and a Subtree Prune and Regraft (SPR) radius of 3. The second run used the output of reconciled trees from the first run as input, and switched to an SPR radius of 5, and the Q.PFAM amino acid substitution model (83), which was trained on the Pfam dataset. We did not use NQ.PFAM, because time non-reversible models are only implemented in IQ-Tree (82), and not in GeneRax. In both runs, we used the UndatedDTL probabilistic model to compute the reconciliation likelihood. The second run of GeneRax reduced estimated transfer rates by an additional 7% (Welch two sample t-test, p = 10^-12^), indicating continued improvements to the phylogenies.

We re-estimated the branch lengths of the reconciled Pfam trees in IQ-Tree using the NQ.PFAM substitution model with no rate heterogeneity, then performed midpoint rooting using the phytools R package (84) on these re-estimated branch lengths. As alternative rooting methods, we also explored and rejected minimum variance (85), minimal ancestral deviation (86), and rootstraps based on time non-reversible substitution models (87). The first two methods work best when deviations from the molecular clock average out on longer timescales, which is not true for phylogenies in which evolution e.g. at different temperatures causes sustained differences in evolutionary rate. Indeed, minimum variance failed to resolve the prokaryotic supergroups as separate clades, in visual inspection of PF00001, due to presumed genuine rate variation among taxa. The latter produced very low confidence roots. In contrast, midpoint rooting largely conformed to expectations for aaRSs once we implemented the procedure for outlier removal described under “Classifying Pfam domains into ancient phylostrata” below.

We then implemented a new --enforce-gene-tree-root option in GeneRax, and ran GeneRax in evaluation mode, with Q.PFAM+G as the substitution and rate heterogeneity models, respectively. Evaluation mode re-estimates the reconciliation likelihood and the duplication, transfer and loss (DTL) rates on a fixed tree, without initiating a tree search. Fifteen reconciled Pfam trees had inferred transfer rates higher than 0.6, three times the seed transfer rate implemented by GeneRax. We took this as a sign of poor tree quality, and annotated these 15 Pfams as of unclassifiable age.

### Filtering out HGT between archaea and bacteria

Exclusion of horizontal gene transfer (HGT) between bacteria and archaea facilitates the classification of a Pfam into LUCA (Figure 2a). To achieve this, we divided sequences into “homogeneous groups”, meaning the largest monophyletic group in the Pfam tree for which the corresponding species all belong to the same prokaryotic supergroup. Each homogeneous group was considered as a candidate for exclusion, via its “focal node” separating it from its sister group. To avoid over-pruning, we do not consider deep focal nodes that are 2 or fewer nodes away from the root.

To be excluded, we first require the focal node to be ‘mixed’, meaning its descendants are found within both Bacteria and Archaea. We next require the focal node to be labelled by GeneRax as most likely a transfer (T), rather than a duplication (D) or speciation (S). Finally, to identify homogeneous groups likely to be receivers rather than the donors of transferred sequences, we require the sister lineage to contain no sequences present in the same supergroup as that defining the homogeneous group in question. An example of filtering is shown in Figure 2b.

We ran the filtering process twice to address rare occasions of an intradomain HGT nested within another intradomain HGT group. In the second filter, we apply the third criterion after pruning the homogenous groups identified as HGT during the first filter.

### Classifying Pfam domains into ancient phylostrata

We re-rooted the HGT-pruned Pfam trees using the midpoint.root function in the ‘phytools’ R package (84), before classifying them into phylostrata (i.e. cohort of sequences of similar age). Classification was based on the locations of the most recent common ancestors (MRCAs) of each supergroup. For a LUCA Pfam, we require the root to separate the MRCAs of all bacterial supergroups from the MRCAs of all archaeal supergroups (Figure 2a).

If there were no horizontal transfer, and the tree of a Pfam present in one copy in LUCA were error-free, then the MRCAs for the nine supergroups would be two to four branches away from the root. This is true even if our Pfam tree and/or species tree do not correctly capture the true phylogenetic relationships among supergroups. However, we cannot ignore HGT; we did not filter out the products of HGT between supergroups within Archaea or within Bacteria, only that of HGT between Archaea and Bacteria. HGT from a more derived supergroup to a more basal supergroup will move the inferred MRCA of the former further back in time. Given rampant HGT, whether real or erroneously implied by Pfam tree error, we required Pfams to have their supergroups’ MRCA two branches away from the root (Figure 2a).

Phylogenies with three or more basal bacterial supergroups and two or more basal archaeal supergroups were classified as LUCA. In other words, we allow the absence of up to two supergroups per taxonomic domain, as compatible with ancestral presence followed by subsequent loss. Trees with three or more basal bacterial supergroups but fewer than two basal archaeal supergroups, as well as trees with two or more basal archaeal supergroups but fewer than three basal bacterial supergroups, were classified as ancient but post-LUCA. These are candidate Pfams for the Last Bacterial Common Ancestor (LBCA) and the Last Archaeal Common Ancestor (LACA) phylostrata, respectively, but the necessary HGT filtering for sufficient confidence in this classification is beyond the scope of the current work. If only one basal supergroup is present, then the Pfam is classified into the corresponding supergroup-specific phylostratum, meaning it emerged relatively recently (modern post-LUCA). If two basal bacterial supergroups (and no archaeal supergroups) were present, the Pfam was classified as post-LBCA which was also considered modern post-LUCA (younger than LBCA but older than the supergroup-specific phylostrata). The remaining Pfams were considered unclassifiable.

We also classify into a pre-LUCA phylostratum the subset of LUCA-classified Pfams for which there is evidence that LUCA contained at least two copies that left distinct descendants. This is motivated by the assumption that LUCA domains that were born earlier are more likely to have duplicated and diverged prior to the archaeal-bacterial split (88). We require that both the nodes that are only one branch from the root be classified as LUCA nodes. This means that each of these nodes should, after HGT filtering: i) split a pure-bacterial lineage from a pure-archaeal lineage, and ii) include as descendants at least three bacterial and two archaeal basal MRCAs no more than two nodes downstream of the potential LUCA nodes (Figure 2c).

Assignment of a Pfam to a phylostratum is sensitive to the root’s position. Midpoint rooting is based on the longest distance between two extant sequences. A single inaccurately placed sequence can yield an abnormally long terminal branch, upon which the root is then based. This phenomenon was readily apparent upon manual inspection of rooted Pfam trees. To ensure the robustness of age classifications to the occasional misplaced sequence, we removed the Pfam instance with the longest root-to-tip branch length in each HGT-filtered tree as potentially faulty, re-calculated the midpoint root, and then re-classified each Pfam. We repeated this for ten iterations, then retained only those Pfams that were classified into the same phylostratum at least 7 out of 10 times. Our HGT filtering algorithm does not act on nodes near the root, making it robust to small differences in root position; we therefore did not repeat the HGT-filtering during these iterations.

We classified clans that contained at least two LUCA Pfams as pre-LUCA clans. Clans that contained both ancient archaeal and ancient bacterial post-LUCA Pfams (i.e. candidate LACA and LBCA Pfams) were classified as LUCA. Clans that contained at least two different archaeal but no bacterial supergroup-specific Pfams, or three different bacterial supergroup-specific Pfams but no archaeal supergroup-specific Pfams, were classified as ancient post-LUCA clans. Clans that meet neither of these criteria, and that contain at least one unclassified Pfam, were considered unclassifiable due to the possibility that the unclassified Pfam might be older than the classified Pfams present in the clan. All other clans were assigned the age of their oldest Pfam.

For a more stringent analysis of amino acid usage, we restrict our Pfam dataset to those present in proteins annotated by Moody et al. (21) as >75% likely to be in LUCA. We then re-classified clan ages. Data on the likelihood of Pfams being present in LUCA, as annotated by Moody et al. (21), can be found in ‘MoodyPfams_probabilities.csv’ on GitHub.

### Ancestral amino acid usages

Ancestral sequence reconstruction (ASR) can introduce a variety of biases. ASR methods do not resolve alignment gaps well, to infer indel evolution, instead inferring ancestral sequences far longer than any contemporary descendant. To avoid bias among amino acids regarding which contemporary sequences appear in the ancestral sequence more often than they should, we retain only sites where more than 50% of the sequences contain an amino acid (i.e. no indel). This ensures that no amino acid can be double counted.

For Pfams classified as pre-LUCA or LUCA, we require that a given site contain an amino acid and not a gap in at least five bacterial sequences and five archaeal sequences. This additional filter helps ensure that the ancestrally reconstructed sites were not inserted post-LUCA (even when the Pfam itself dates back to LUCA). It also reduces the impact of any Pfams misclassified as ancient on the inferred ancient amino acid usage.

Following these filters, we ran the remaining sites in each Pfam alignment (prior to HGT filtering) through IQ-Tree with the -asr option, the NQ.PFAM substitution model, and R10 rate heterogeneity. We then excluded low confidence sites from subsequent analyses, based on the most likely amino acid having an ancestral probability estimate <0.4. Combined with the other two filters described above, the concatenated sequence length for all four phylostrata (pre-LUCA, LUCA, post-LUCA, and modern) fell by ∼11%, presumably preferentially excluding rapidly evolving sites to a similar degree in all four cases, such that amino acid exclusion biases cancel out when ratios are taken.

We then summed over the amino acid probability distributions at each site at the deepest node, and divided by the number of sites, to obtain per-Pfam estimated ancestral amino acid frequencies. For each clan, we took the ancestral amino acid frequencies across Pfams, weighted by the number of ancestral sites in the Pfams. For each phylostratum, we averaged across clans, weighted by the maximum number of ancestral sites across all Pfams in a given clan. We calculated a standard error associated with each phylostratum mean using the weighted_se() function in the diagis R package (89).

We divided ancestral amino acid frequencies for the LUCA and pre-LUCA phylostrata by post-LUCA ancestral amino acid frequencies to produce measures of relative usage. Standard errors of each of these ratios *L/P* were calculated using an approximation derived from a Taylor expansion of the ratio: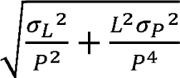(90). These were used in weighted linear model 1 regressions, using the lm() function with the ‘weights’ argument in the ‘stats’ package in base R (91). Uncertainty in the ancestral states arising over 4 billion years of evolution is expected to bring values of *L/P* closer to one, without entirely erasing the signal. As a negative control for bias, we calculate the relative amino acid usage of post-LUCA clans by dividing the ancestral amino acid frequencies for post-LUCA clans by the ancestral amino acid frequencies for modern clans.

Standard errors in Trifonov’s (4) average rank reflect but underestimate uncertainty; we therefore treat Trifonov’s (4) rankings as the dependent variable and use its weights rather than errors on to *L/P* weight the regression model in Figure 4c. Standard errors are not available for alternative results based on Trifonov’s 2004 order (38).

### Hydrophobic interspersion

The degree to which hydrophobic are clustered vs. interspersed along the primary sequence was calculated as a normalized index of dispersion for each Pfam instance (35). This metric uses the ratio of the variance to the mean in the number of the most hydrophobic amino acids (leucine, isoleucine, valine, phenylalanine, methionine, and tryptophan) within consecutive blocks of six amino acids. The values of this index of dispersion were then normalized, to make them comparable across Pfams with different lengths and hydrophobicities. In cases where the Pfam length was not a multiple of 6, the average across all possible 6-amino acid frames was computed, trimming the ends as needed. For additional details, see Foy et al. (36) or James et al. (14). For each Pfam, we then took the average across all its instances (prior to downsampling species).

### Transmembrane annotation

We identified transmembrane sites within each Pfam using DeepTMHMM (92) on a consensus sequence generated from the original multiple sequence alignments (prior to HGT filtering) using the majority-rule seq_consensus() function in the R package ‘bioseq’ (93).

## Data and Code Availability

Data files and R scripts used to generate the results and figures are available at sawsanwehbi/Pfam-age-classification GitHub repository. Pfam sequences, alignments, trees and mappings to protein IDs are available at https://figshare.com/projects/Pfam-age-classification-data/201630.

## Acknowledgments

We thank NASA [80NSSC24K0384] and the John Templeton Foundation [62220] for funding SW, NM, and JM, Chan-Zuckerberg Initiative [EOSS4-0000000312] for funding BQM, the DFG [STA 860/6-2] for funding BM, the NSF [DEB-2333243] for funding AW, NM, and JM, the Arizona NASA Space Grant Consortium [80NSSC20M0041] for funding NM, and the NIH [T32GM132008] for funding AW. We thank Mike Barker, Alan Moses, and Elisa Tomat for helpful discussions, and Cat Wolner for comments on the manuscript.

