## Supplemental Figures and Tables for "Order of amino acid recruitment into the genetic code resolved by Last Universal Common Ancestor’s protein domains"

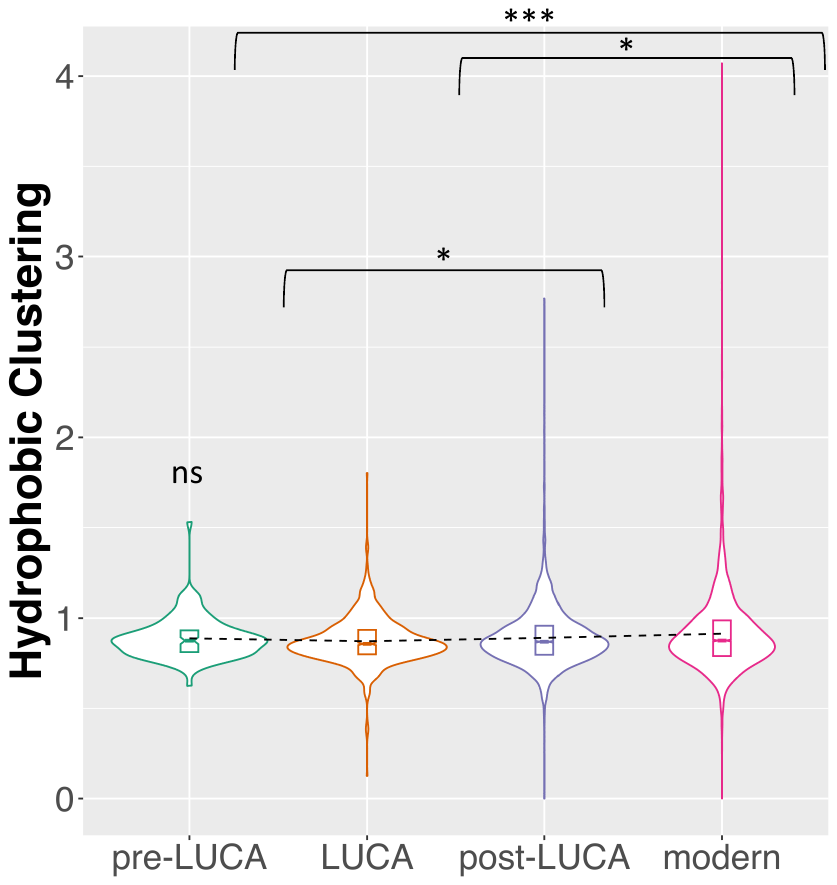

**Supplementary Figure 1. Older protein Pfams have more interspersed hydrophobic amino acids.** Independent amino acid placements correspond to a “hydrophobic clustering” score of 1, values less than 1 indicate interspersed hydrophobic amino acids along the primary sequence, values greater than 1 indicate clustered hydrophobic amino acids; see Methods for details. Hydrophobic amino acids are more interspersed (with lower clustering value) in LUCA Pfams than in either post-LUCA Pfams (p = 0.02) or modern Pfams (p = 0.0003). The latter two also differ from each other (p = 0.02). Pairwise Wilcoxon rank sum tests were used to test for statistical significance. Boxplots show the median, upper, and lower quartiles of the data. Violin plots were generated using the geom_violin() function in the ggplot R package (1). Small notches around the medians define the 95% confidence intervals for the medians, and dashed lines join means. The hydrophobic clustering of pre-LUCA Pfams is not significantly different from any of the other phylostrata, which may be due to the low number of per-LUCA Pfams. We note that unlike the animal proteins analyzed in James et al. (2), even the youngest, modern Pfams have clustering significantly below the expected value of 1 for independent sites (Binomial test; p < 10^-16^).

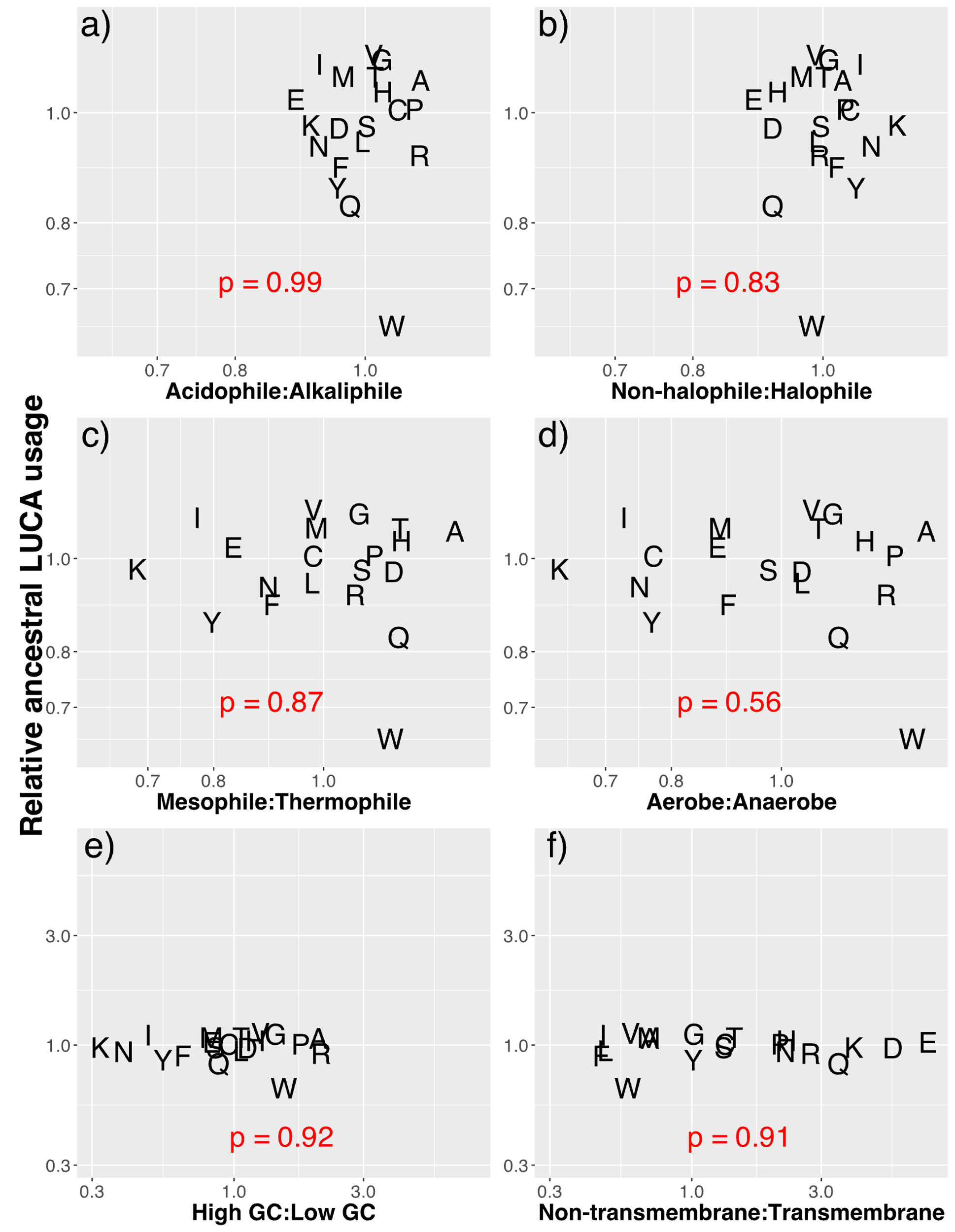

**Supplementary Figure 2. Environmental, genomic and structural amino acid biases do not correlate with the ancestral amino acid usage in LUCA clans**. We compare relative amino acid usage in different conditions (x-axes) to that of our inferred ancient amino acid usage in LUCA clans (y-axis). Both axes are plotted on a natural logarithmic scale with back transformed tick labels. Different scales on the bottom row help show how LUCA enrichments are of larger magnitude than some factors (a-b), similar to some (c-d), and smaller than others (e-f). a-c) We calculated the proteomic amino acid usage as a function of environmental factors, using the proteomic and environmental data published by Amangeldina et al.(3). d) The oxygen tolerance of the prokaryotic species in the Amangeldina et al. (3) dataset was taken from the BacDive database (4). e) We compared the amino acid usage of organisms with > 60% GC to those with < 40% GC using GC content data from 11,710 representative prokaryotic genomes from the NCBI Refseq Prokaryotic Genomes database, taken from <https://chrisgaby.github.io/post/prokaryotic-genome-size/index.html>) (5) (accessed August 8^th^ , 2024) . f) 1179 transmembrane Pfams (out of 8282 total Pfams) were identified based on the predicted presence of transmembrane alpha helices or beta sheets using DeepTMHMM (6). We compared the amino acid frequencies of non-transmembrane Pfams to those in transmembrane regions within transmembrane Pfams. Associated data can be found in ‘Environment_AAC.csv’, ‘Bacdive_Oxygen_requirement.csv’, ‘prokaryotes_GC.csv’ and ‘DEEPTmhmm_consensusPfam.out’ files on GitHub.

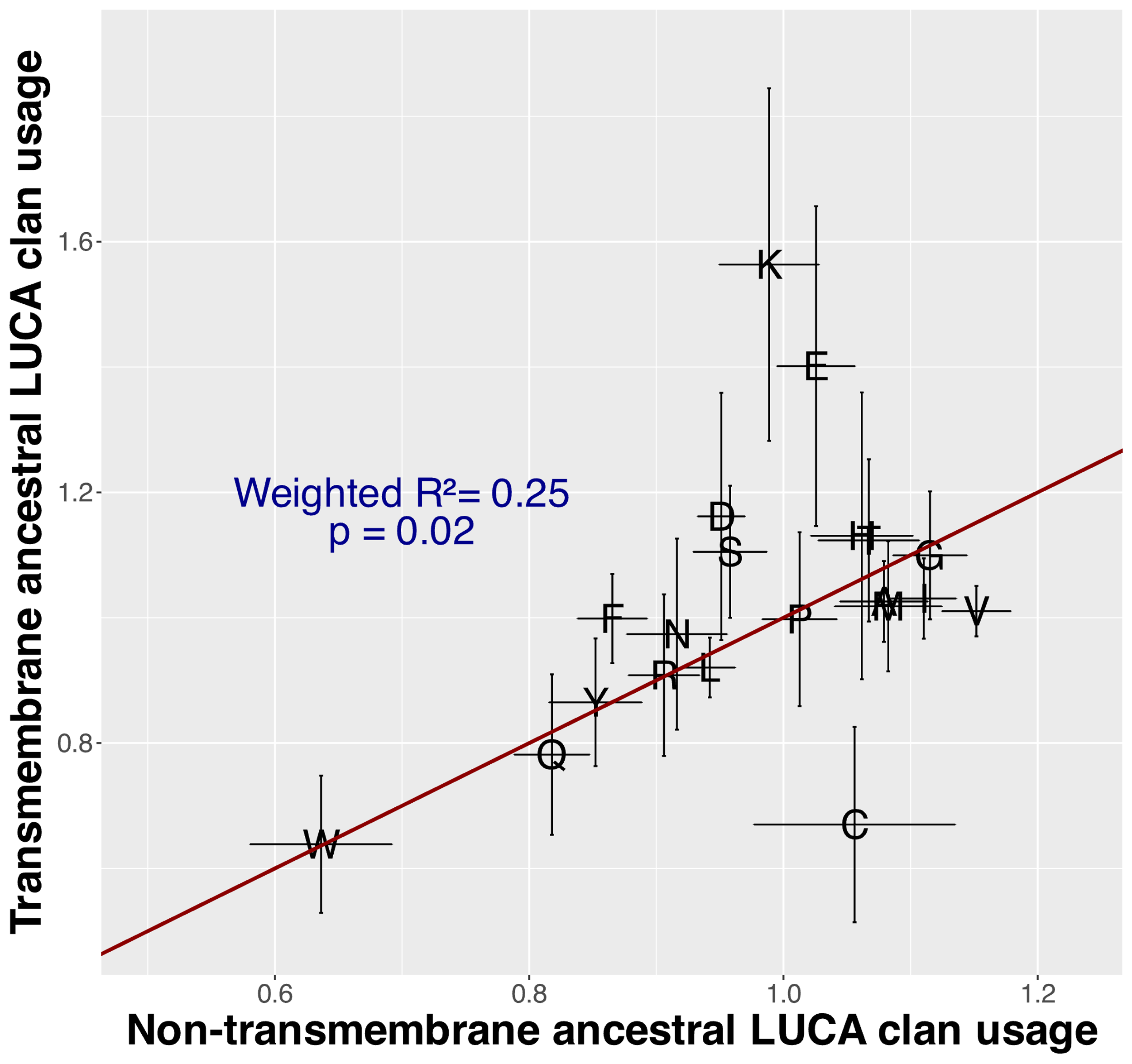

**Supplementary Figure 3. Ancient amino acid usage at transmembrane sites resembles that of non-transmembrane clans.** LUCA and post-LUCA clans were used only if they contain either exclusively transmembrane or exclusively non-transmembrane Pfams. We calculated transmembrane amino acid frequencies only at transmembrane sites as predicted by DeepTMHMM (6). Error bars represent the standard errors of each LUCA / post-LUCA ratio, approximated via a Taylor expansion (7); see Methods. The R^2^ and p value are estimated from a weighted model 1 regression, using the larger errors for transmembrane usage and neglecting errors in non-transmembrane usage. The y=x line is shown in red.

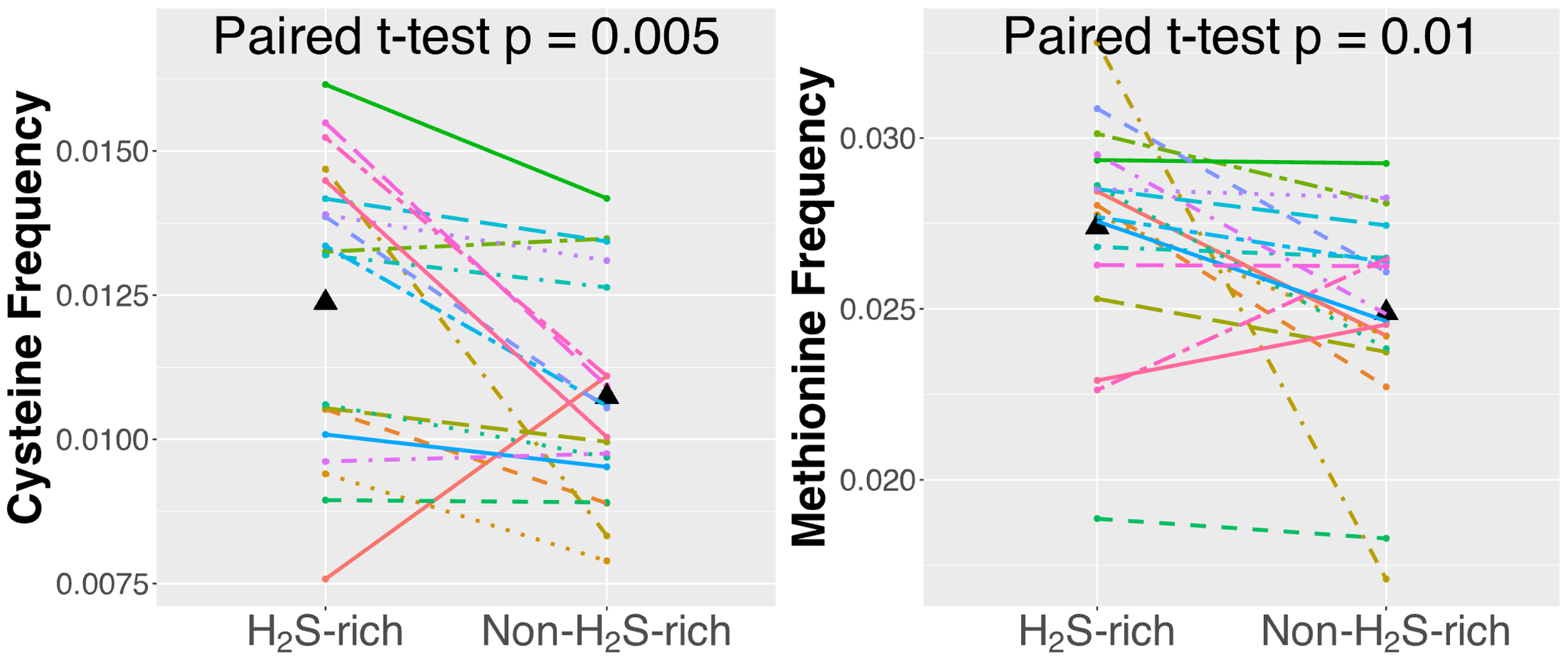

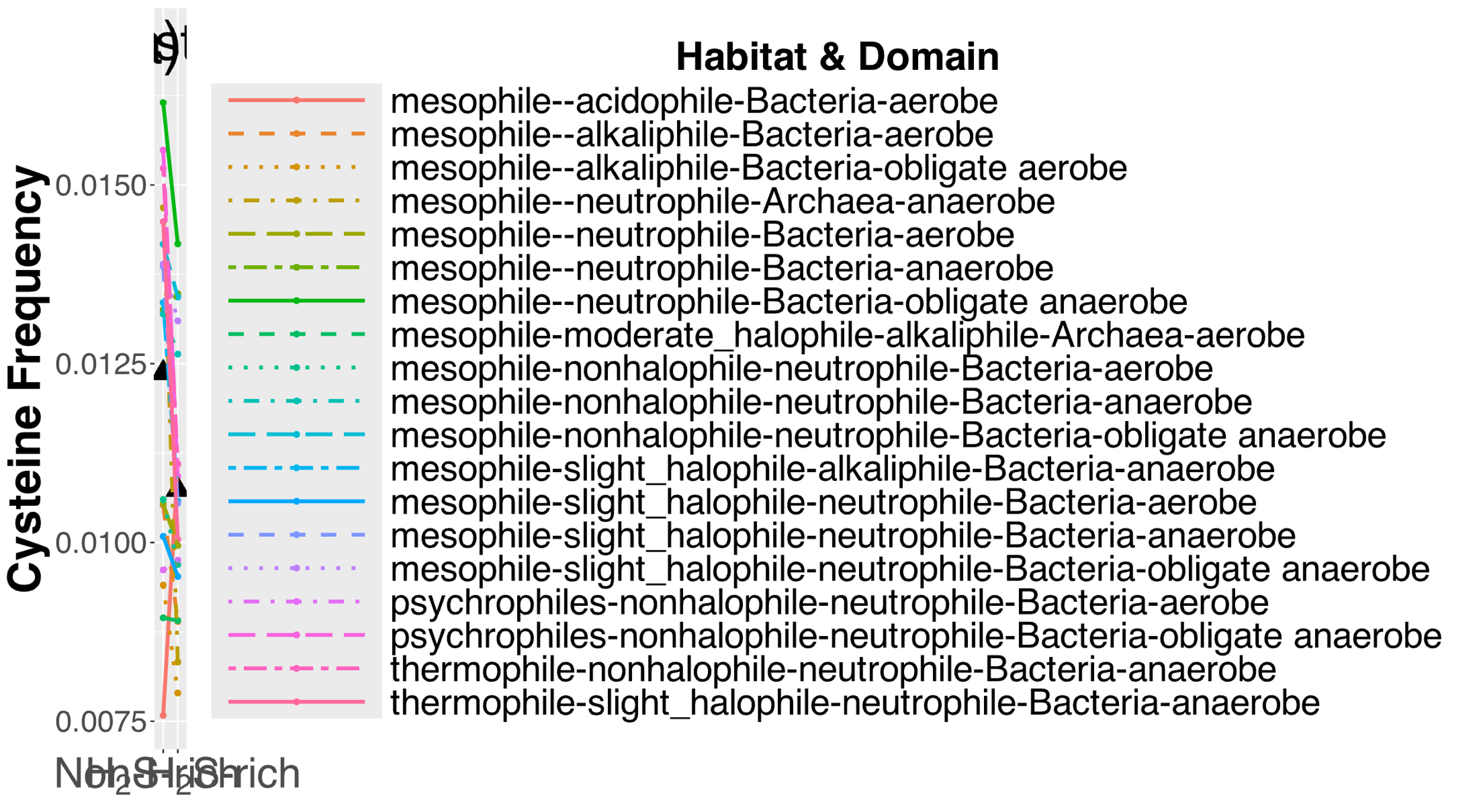

**Supplementary Figure 4. Sulfur-containing amino acids, M and C, are used more by contemporary prokaryotes living in H_2_S-rich environments.** Nineteen colored points and connected lines indicate matched pairs representing a range of environments (temperature, salinity, pH, and oxygen) and both Bacteria and Archaea. Triangles indicate means. We took data on habitat, taxonomy and amino acid frequencies from Amangeldina et al. (3). Each data point averages across a set of genera that inhabit similar environments and belong to the same taxonomic domain. One species was randomly sampled per genus to avoid pseudo-replication. Each line then joins the averages between the two sets of genera that inhabit otherwise similar environments and belong to the same taxonomic domain. Organisms living in H_2_S-rich environments include sulfur-reducing and sulfur-oxidizing prokaryotes. A list of the species used in this analysis, with their respective environmental characteristics, and M and C frequencies can be found in ‘sulfur_nonsulfur_species.csv’ on GitHub.

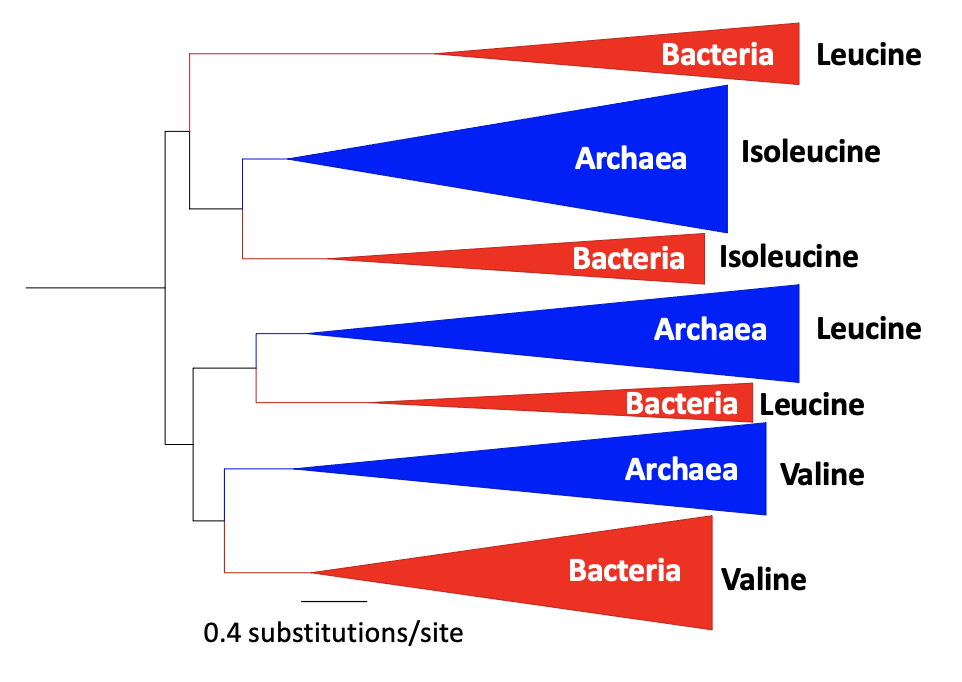

**Supplementary Figure 5. Topology of the leucine, valine, and isoleucine tRNA synthetase core catalytic domain confirms pre-LUCA origin.** Three archaeal-bacterial splits denoting LUCA are shown, representing leucine, valine and isoleucine specific tRNA synthetase homologs. Collapsed archaeal clades names are in blue and collapsed bacterial clades are in red. Branch lengths were estimated using the Pfam-trained time non-reversible amino acid substitution model NQ.pfam (8), with an R10 heterogeneity model. Tree depicted is HGT-filtered, and midpoint rooted following pruning of the first ten branch length outliers. **Supplementary Table 1.** Pfams with classified ages, ancestral amino acid frequencies, conserved sequence length, hydrophobic clustering score and clan data are also found in ‘Pfam_data_ancestralAAC.csv’ at [sawsanwehbi/Pfam-age-classification GitHub repository](https://github.com/sawsanwehbi/Pfam-age-classification/tree/main).

**Supplementary Table 2.** Clans with classified ages, ancestral amino acid frequencies and maximum conserved sequence length are also found in ‘Clan_data_ancestralAAC.csv’ at [sawsanwehbi/Pfam-age-classification GitHub repository](https://github.com/sawsanwehbi/Pfam-age-classification/tree/main).

**Supplementary Table 3. Our Pfam classifications confirm previous inference of LUCA’s metabolism.** All but two Pfams associated with the enzymes in the hydrogen (H_2_) metabolism, assimilatory nitrate reduction, assimilatory sulfate reduction and Wood-Ljungdahl pathway were present in LUCA. Pfams associated with the nitrogenase family were not present in LUCA.

| PATHWAY | KEGG ID | GENE NAME | PFAM IN LUCA | PFAM NOT IN LUCA |
| --- | --- | --- | --- | --- |
| Dihydrogen  Metabolism | K06281 | hydrogenase large subunit | PF00374 |  |
|  | K14126 | F420-non-reducing hydrogenase large subunit | PF00375 |  |
| Assimilatory  Nitrate Reduction | K00367 | ferredoxin-nitrate reductase | PF04879;PF00384  PF01568 |  |
| Assimilatory  Sulfate Reduction | K00957 | sulfate adenylyltransferase | PF01507 |  |
|  | K00392 | sulfite reductase | PF01077;PF03460 |  |
|  | K00930 | phosphoadenosine phosphosulfate reductase | PF00696 |  |
| Wood-Ljungdhal  Pathway | K00198 | carbon-monoxide dehydrogenase catalytic subunit | PF03063 |  |
|  | K05299 | formate dehydrogenase (NADP+) alpha subunit | PF13510;PF12838  PF04879;PF00384 |  |
|  | K15022 | formate dehydrogenase (NADP+) beta subunit | PF14691;PF12838  PF04060;PF07992 | PF14691 (LBCA) |
|  | K22015 | formate dehydrogenase (hydrogenase) | PF04879;PF00384  PF01568 |  |
|  | K25124 | FeS-containing electron transfer protein | PF13247;PF12800 |  |
|  | K01491 | methylenetetrahydrofolate dehydrogenase (NADP+) | PF02882 | PF00763 (unclassifiable) |
|  | K01500 | methenyltetrahydrofolate cyclohydrolase | PF04961 |  |
|  | K00297 | methylenetetrahydrofolate reductase (NADH) | PF02219 |  |
|  | K25007 | methylenetetrahydrofolate reductase (NADH) small subunit | PF12225 |  |
|  | K15023 | 5-methyltetrahydrofolate corrinoid/iron sulfur protein methyltransferase | PF00809 |  |
|  | K00197 | acetyl-CoA decarbonylase/synthase, CODH/ACS complex subunit gamma | PF03599;PF04060 |  |
|  | K00194 | acetyl-CoA decarbonylase/synthase, CODH/ACS complex subunit delta | PF03599 |  |
| Nitrogen  Fixation | K02586 | nitrogenase molybdenum-iron protein alpha chain |  | PF00148 (LBCA) |
|  | K02587 | nitrogenase molybdenum-cofactor synthesis protein NifE |  | PF00148 (LBCA) |
|  | K02592 | nitrogenase molybdenum-iron protein NifN |  | PF00148 (LBCA) |
|  | K02588 | nitrogenase iron protein NifH |  | PF00142 (unclassifiable) |
|  | K02591 | nitrogenase molybdenum-iron protein beta chain |  | PF00148 (LBCA);PF11844 (Proteobacteria) |
|  | K02593 | nitrogen fixation protein NifT |  | PF06988 (Proteobacteria) |
